# Guillain-Barre syndrome outbreak in Peru: Association with polymorphisms in *IL-17, ICAM-1* and *CD1*

**DOI:** 10.1101/667154

**Authors:** Luis Jaramillo-Valverde, Kelly S Levano, Isolina Villanueva, Meylin Hidalgo, Marco Cornejo, Pilar Mazzetti, Mario Cornejo-Olivas, Cesar Sanchez, Julio A Poterico, Julio Valdivia-Silva, Heinner Guio

## Abstract

Guillain-Barre Syndrome (GBS) is considered a complex disorder with significant environmental effect and genetic susceptibility. Genetic polymorphisms in *CD1E, CD1A, IL-17* and/or *ICAM-1* genes had been proposed as susceptibility genetic variants for GBS mainly in Caucasian population. This study explores the association between selected polymorphisms in these genes and GBS susceptibility in confirmed GBS cases reported in mestizo population from northern Peru during the most recent GBS outbreak of May 2018. A total of 9 non-related cases and 11 controls were sequenced for the polymorphic regions of *CD1A, CD1E, IL-17* and *ICAM-1* genes. We found a significant protective association between heterozygous GA genotype in *ICAM-1* gene (241Gly / Arg) and GBS (p <0.047). *IL-17* was monomorphic in both controls and patients. No significant differences were found in the frequency of SNPs in *CD1A* and *CD1E* between the group with GBS patients and healthy controls. Further studies with larger sample size will be required to validate these findings.

## INTRODUCTION

Guillain-Barre syndrome (GBS) is an acute inflammatory polyradiculoneuropathy with ascending weakness staring in lower limbs, extension to the upper limbs and face, as well as complete loss of deep tendon reflexes (1-4). The annual incidence of GBS is 0.5-2 cases per 100,000 people, which increases with age (3,4). GBS is rare in children under two years (5); males are 1.5 times more likely to suffer GBS than women (4-6). The exact cause of GBS has not been defined yet; however, 50-70% of the cases appear 1-2 weeks after an infection (bacterial or viral) inducing an aberrant autoimmune response directed to the peripheral nerves and their spinal roots (4,7,8). GBS is known to occur in several forms including acute inflammatory demyelinating polyradiculoneuropathy (AIDP), Miller Fisher syndrome (MFS), acute motor axonal neuropathy (AMAN) and acute motor-sensory axonal neuropathy (AMSAN).

The interaction between microbial and host factors has been poorly studied in GBS, as well as the genetic susceptibility of an individual to develop this syndrome. Possible markers of genetic susceptibility to GBS have been reported, including *CD1E* (OMIM #188411), *CD1A* (OMIM #188370*), IL-17* (OMIM #606496) and *ICAM-1* (OMIM #147840) (9,10). The *CD1E* and *CD1A* genes are glycoproteins of Major Histocompatibility Complex (MHC) specialized in capturing and presenting glycolipids to T cells (10,11). In a research study conducted by Caporale et al, it was reported that individuals with the genotype *CD1E*\*01/01 were 2.5 times more susceptible to develop GBS, while individuals with the genotypes *CD1A*\*01/02 or *CD1E*\*01/02 had a risk of 3.6 and 2.3 times lower, respectively (10). Likewise, there has also been an association of GBS with polymorphisms *IL-17* (Glu126Gly) and *ICAM-1* (Gly241Arg) (9). Il-17 regulates the expression of inflammatory genes, including proinflammatory chemokines, hematopoietic cytokines, acute phase response genes and antimicrobials (12) in neutrophils, macrophages and endothelial cells (13). On the other hand, previous studies show that *ICAM-1* plays a central role in the development of demyelinating disease (14).

GBS and its association with a variety of infectious agents have been reported in Peruvian population. By 2014, case series of 32 GBS cases followed in Lima (capital city) found that AIDP was the most common form (75%) followed by AMAN and MFS with frequencies of 18.8% and 6.3%, respectively (15). By contrast, series from northern Peru (2017) found 16 Peruvian cases where AMSAN was the most common form (37.5%) followed by AMAN (25%) and AIDP (12.5%) (16). In 1987, five GBS cases were associated with a viral infection caused by a rabies vaccine prepared with the brain of a lactating mouse (17). In 2010, a GBS case was reported associated to Brucellosis, an infectious disease caused by Brucella bacteria genus (18).

In Peru, 15 cases of GBS were reported between April and May of 2018 in Trujillo, northern Peru, during summer time, activating a national epidemiological alert declared by the Ministry of Health. All cases were put on immunoglobulin G and managed in the intensive care unit at a regional Hospital. Blood samples were taken in all cases for both environmental exposure and DNA extraction for further genetic analysis.

This study determines the occurrence of polymorphisms in *IL-17, ICAM-1* and *CD1* genes in GBS cases with a medical history of enteric respiratory and / or gastrointestinal infection and controls.

## PATIENTS AND METHODS

### Ethical approval

This study was approved by the ethics and research committee of Belen Hospital of Trujillo, northern Peru. A written informed consent was obtained from all subjects prior to recruitment for the study.

### Cases and controls

Nine patients with GBS (7 men and 2 women, age: 52 to 65 years) followed at a regional hospital in northern Peru were enrolled in the study during the outbreak of GBS occurred in May, 2018. Eleven healthy subjects (7 women and 4 men, age: 27 to 74 years) were randomly selected as controls from the same geographical area of residence. A total of 3mL of blood was obtained from peripheral veins in all subjects.

### Isolation of DNA and genotyping of IL-17, ICAM-1 and CD1 genes

100 microliters of peripheral blood were used to extract DNA using QIAamp DNA Blood Kit Mini (Qiagen, CA, USA) and INBIOMag Genomic DNA Kit (INBIOMEDIC, Peru). According to the literature and genotypes location reported, the following fragments were selected: *IL-17, ICAM-1* and *CD1* (2 fragments, found in exon 2). Specific primers were designed for each DNA fragment (Table 1) and the fragments were PCR amplified using Taq PCR Master Mix Kit (Qiagen, CA, USA). PCR products were purified and sequenced by Sanger technique in Macrogen (Soul Korea).

**Table 1.**
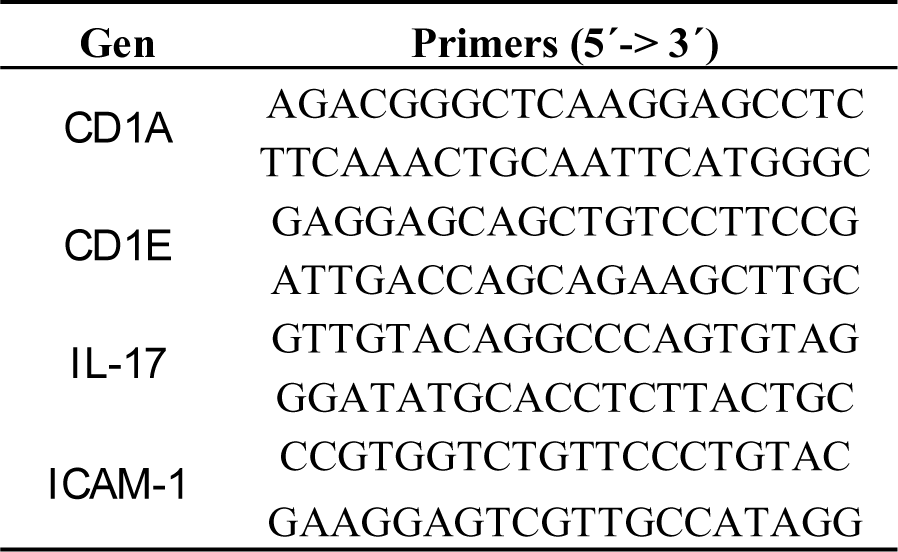
Primers used in the genetic analysis of Guillain-Barre Syndrome

### Genetic analysis

The DNA sequence data was processed using Geneious R11 software (Biomatters Ltd.). The polymorphisms *IL-17* (Glu126Gly), *ICAM-1* (Gly241Arg) and *CD1A* and *CD1E* genotypes were evaluated. The sequences obtained from *IL-17, ICAM-1* and *CD1* were compared with sequences reported in previous research and/or global database.

### Statistical analysis

Polimorfisms of *IL-17, ICAM-1 and CD1* were listed by frequency and percentage. Exploratory analysis comparing frequency of polymorphisms between cases and controls were performed by the chi square test and logistic regression models. Results were considered significant if p <0.05. Statistical analysis were performed by SATA version 15.0 (Illinois, USA).

## RESULTS

A total of 9 cases (7 males, 77.8%) analyzed (Table 2) were diagnosed as SGB of atypical presentation. Eight patients reported some type of symptomatology 8 weeks before the onset of paralysis such as respiratory, gastrointestinal infections, non-purulent conjunctivitis, joint and head pain. Only one reported a trip to Virú province days before the paralysis.

**Table 2.**
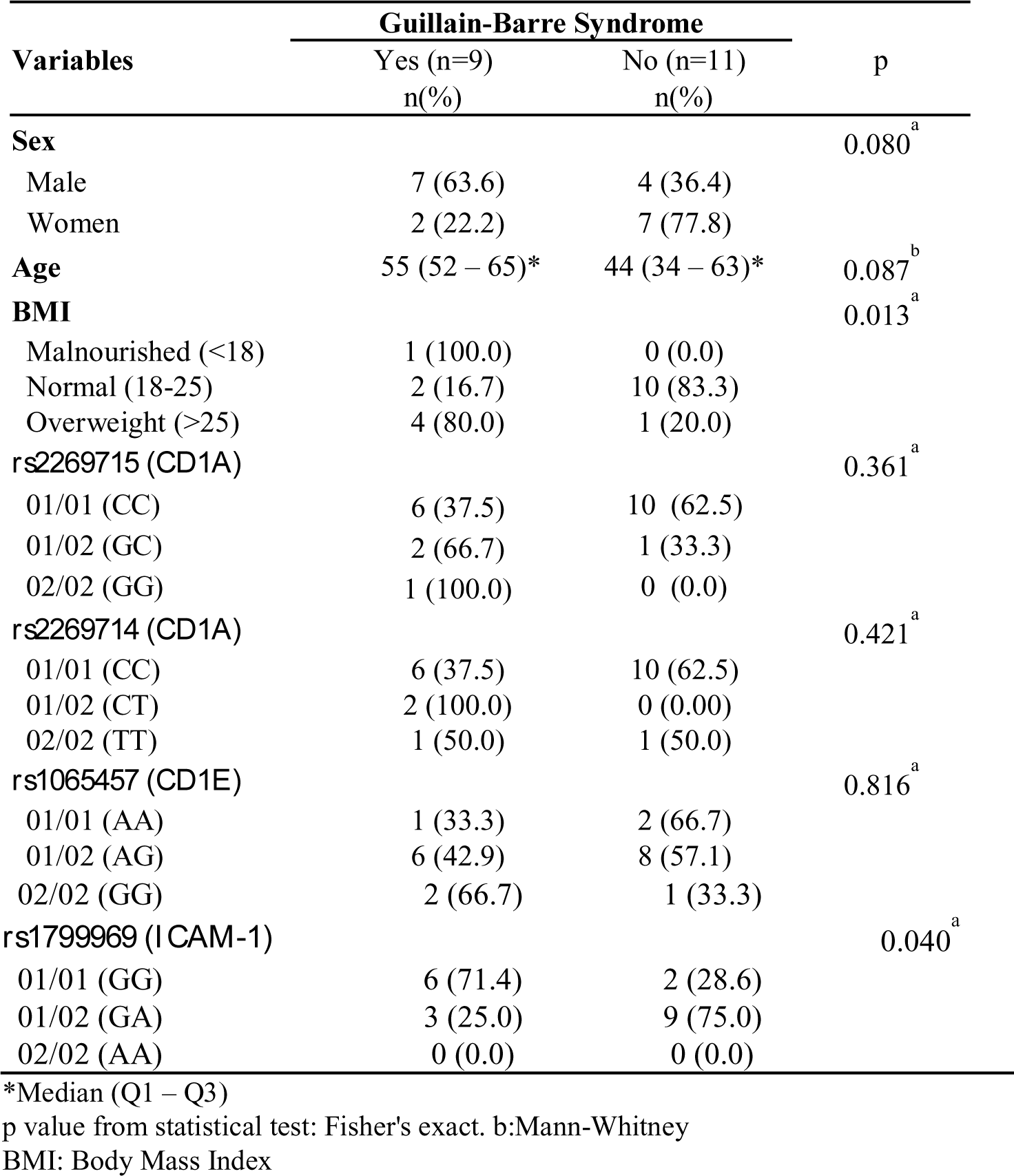
Bivariate analysis of factors associated with the diagnosis of Guillain-Barre Syndrome

All 9 cases experienced muscle weakness, 6 of them also complained of pain, 3 cases presented ataxia, 2 cases had cranial nerves compromise and 3 cases presented symmetric paralysis. Autonomic disturbances, urinary dysfunction, sinus tachycardia and arrhythmia, were each-one demonstrated in one case.

Among controls and GBS cases, *IL-17* gene is monomorphic in 01/01 genotype. Table 3 shows the frequencies of *CD1A, CD1E* and *ICAM-1* alleles and genotypes in controls and patients with GBS. *CD1A* gene is biallelic. Allele 01 is more frequent in both controls and patients with GBS. *CD1A*\*02/02 genotype is not represented in controls and is present in only one of 9 patients with GBS. *CD1A*\*01/01 genotype is slightly more frequent in control patients compared with GBS.

**Table 3:** Genotype and frequency of alleles for *CD1A, CD1E, IL-17* and *ICAM-1* in GBS patients and controls

|  | Genotype |  |  | % of allele |  | % of persons |  |
| --- | --- | --- | --- | --- | --- | --- | --- |
|  | Persons number (%) |  |  | Frequency |  | Positive for alleles |  |
|  | 01/01 | 01/02 | 02/02 | 01 | 02 | 01 | 02 |
| ***CD1A*** |  |  |  |  |  |  |  |
| *rs2269715* |  |  |  |  |  |  |  |
| GBS | 6 (67%) | 2 (22%) | 1 (11%) | 82 | 18 | 89 | 33 |
| Control | 10 (91%) | 1 (9%) | 0 (0%) | 95 | 5 | 100 | 9 |
| *rs2269714* |  |  |  |  |  |  |  |
| GBS | 6 (67%) | 2 (22%) | 1 (11%) | 82 | 18 | 89 | 33 |
| Control | 10 (91%) | 0 (0%) | 1 (9%) | 91 | 9 | 91 | 9 |
| ***CD1E*** |  |  |  |  |  |  |  |
| GBS | 1 (11%) | 6 (67%) | 2 (22%) | 44 | 56 | 78 | 89 |
| Control | 2 (18%) | 8 (73%) | 1 (9%) | 55 | 45 | 91 | 82 |
| ***IL-17*** |  |  |  |  |  |  |  |
| GBS | 9 (100%) | - | - | 100 | - | 100 | - |
| Control | 11 (100%) | - | - | 100 | - | 100 | - |
| ***ICAM-1*** |  |  |  |  |  |  |  |
| GBS | 6 (67%) | 3 (33%) | 0 (00%) | 83 | 17 | 100 | 33 |
| Control | 0 (0%) | 2 (18%) | 9 (82%) | 9 | 91 | 18 | 100 |

*CD1E* gene has two alleles with approximately the same frequency in controls and patients. (Table 3). *CD1E*\*01/02 genotype is more frequent in both controls and patients with GBS. *ICAM-1* gene is biallelic (Table 3). Alelle 01 is more frequent in patients with GBS than in controls. *ICAM-1*\*01/01 genotype is not represented in controls and is present in 6 of 9 patients with GBS. *ICAM-1*\*02/02 genotype is not represented in GBS patients and is present in 9 of 11 control patients.

According to a first bivariate analysis (Table 2), the risk of being diagnosed with SGB in people with *ICAM-1* GA genotype is lower compared to people with *ICAM-1* GG genotype and this difference is statistically significant (p: 0.040). No statistically significant differences were found between groups of patients studied according to GBS diagnosis and other covariates analyzed: sex, age, genotypes *CD1A, CD1E, IL-17* (p> 0.05).

According to regression analysis, *ICAM-1* genotype and BMI variables contribute statistically to association under study (Table 4); Thus, the risk (OR) of being diagnosed with GBS in people with *ICAM-1* GA genotype is about one third (33%) compared with people with *ICAM-1* GG genotype (95% CI: 0.11 – 0.99; p: 0.047). The rest of covariates with exception of BMI do not contribute statistically significant to the association under study.

**Table 4.**
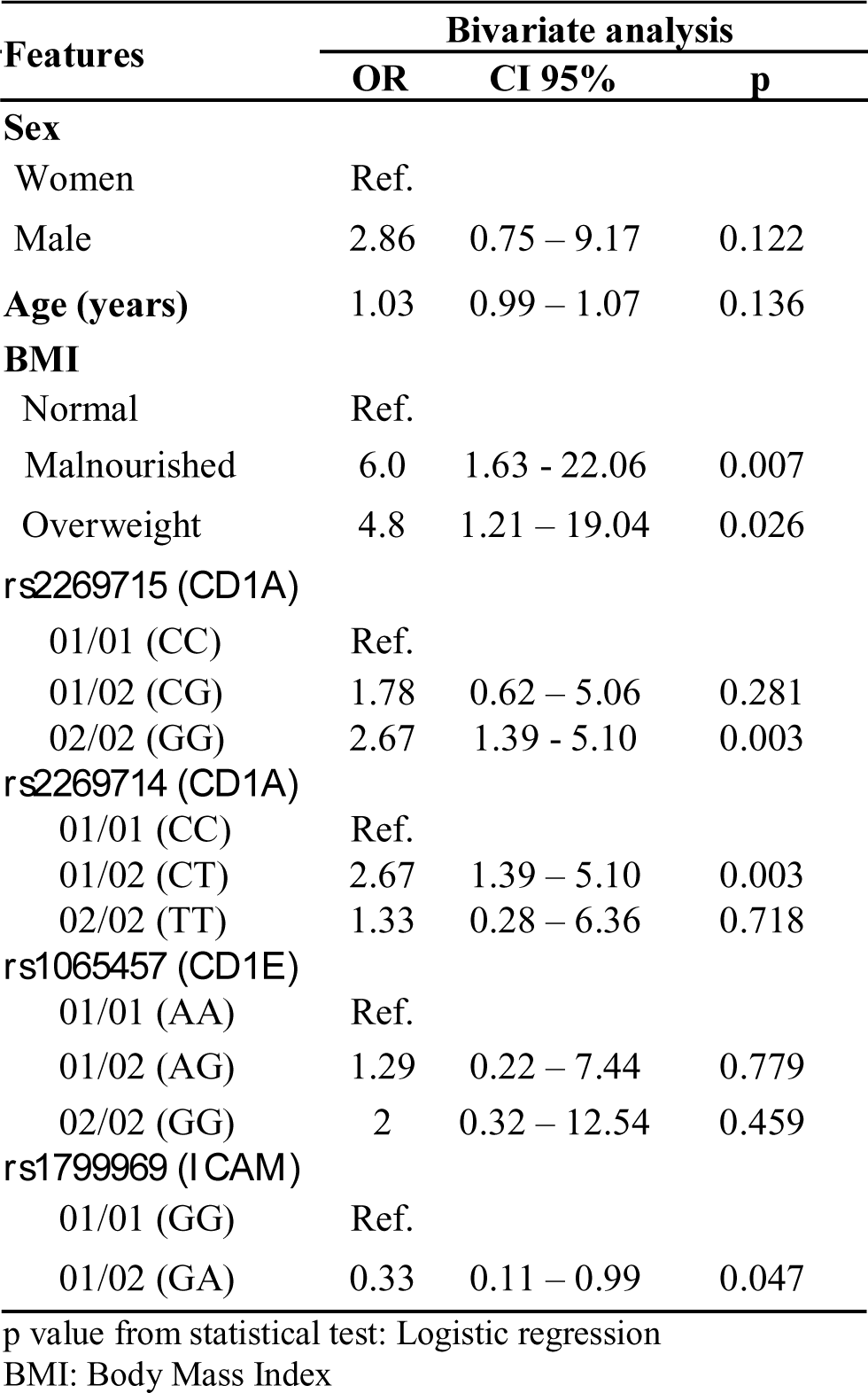
Regression analysis of factors associated with diagnosis of Guillain-Barre Syndrome

| Features | Bivariate analysis |  |  |
| --- | --- | --- | --- |
|  | OR | CI 95% | p |
| **Sex** |  |  |  |
| Women | Ref. |  |  |
| Male | 2.86 | 0.75 – 9.17 | 0.122 |
| **Age (years)** | 1.03 | 0.99 – 1.07 | 0.136 |
| **BMI** |  |  |  |
| Normal | Ref. |  |  |
| Malnourished | 6.0 | 1.63 - 22.06 | 0.007 |
| Overweight | 4.8 | 1.21 – 19.04 | 0.026 |
| ***rs2269715 (CD1A)*** |  |  |  |
| 01/01 (CC) | Ref. |  |  |
| 01/02 (CG) | 1.78 | 0.62 – 5.06 | 0.281 |
| 02/02 (GG) | 2.67 | 1.39 - 5.10 | 0.003 |
| ***rs2269714 (CD1A)*** |  |  |  |
| 01/01 (CC) | Ref. |  |  |
| 01/02 (CT) | 2.67 | 1.39 – 5.10 | 0.003 |
| 02/02 (TT) | 1.33 | 0.28 – 6.36 | 0.718 |
| ***rs1065457 (CD1E)*** |  |  |  |
| 01/01 (AA) | Ref. |  |  |
| 01/02 (AG) | 1.29 | 0.22 – 7.44 | 0.779 |
| 02/02 (GG) | 2 | 0.32 – 12.54 | 0.459 |
| ***rs1799969 (ICAM)*** |  |  |  |
| 01/01 (GG) | Ref. |  |  |
| 01/02 (GA) | 0.33 | 0.11 – 0.99 | 0.047 |

p value from statistical test: Logistic regression BMI: Body Mass Index

## DISCUSSION

This is the first analysis of *IL-17, ICAM-1* and *CD1* genes polymorphisms in Peruvian patients with GBS and healthy controls. Significant differences in the frequency of *ICAM-1* SNPs were observed between patients with GBS and healthy controls, implying *ICAM-1* polymorphisms do influence susceptibility to GBS in the Peruvian population. Additionally, no genetic associations were observed between *IL-17* and *CD1* polymorphisms and GBS susceptibility.

Members of the CD1 family are key players in the immune response to glycolipids and may be involved in the GBS pathogenesis, especially in patients with history of *C. jejuni* infections and anti-ganglioside antibodies (10,19). SNPs in *CD1B* (OMIM #188360), *CD1C* (OMIM #188340), and *CD1D* (OMIM #188410) genes were not determined in the current study because these are very rare and/or silent (21).

*CD1A* and *CD1E* are biallelic in exon 2 (20). *CD1E* is the most polymorphic gene and reports variants in exon 3 also (20, 21). In our study, no significant differences were found in the SNPs frequency of *CD1A* and *CD1E* between GBS patients and healthy controls, which indicates that these genetic polymorphisms do not influence the susceptibility to GBS development in the population studied. (Table 3). In addition, there was no genetic association with clinical outcome in GBS patients. These results do not support the hypothesis that *CD1A* and *CD1E* influence GBS risk as it was raised in a previous study that was based on an Italian cohort of GBS patients (10). It is likely that this discrepancy is caused by differences in patient populations, although both had an ethnic origin, with a similar distribution of polymorphic frequencies in *CD1E* gene, but different in *CD1A* gene. On the other hand, it should be noted that the absence of association with polymorphisms of *CD1* gene does not exclude the possibility that *CD1* molecules play an important role in GBS pathogenesis. More research is needed to determine if it is *CD1* molecules or pathways subsequent to *CD1*, those that participate in process of glycolipids antigenic presentation in GBS.

Many association studies have reported that *IL-17* polymorphisms predispose to autoimmune and inflammatory diseases (22,23,24,25). However, some *IL-17F* polymorphisms (Glu126Gly and His161Arg) may not be significantly associated with autoimmune diseases (26). There are no data reported in the context of *IL-17F* polymorphism with GBS. The importance of *IL-17F* polymorphism in GBS is still largely unknown. In our study, it was observed that *IL-17* gene is monomorphic in 01/01 genotype for patients with GBS and controls.

Increased expression of *ICAM-1* gene has been demonstrated in endothelial cells, microglia and astrocytes in patients with multiple sclerosis (27, 28). We have observed significant protection association with GBS in people with heterozygous GA genotype in *ICAM-1* gene (241Gly / Arg) (p <0.047). These results do not support the association hypothesis of significant risk of heterozygous *ICAM-1* genotype (241Gly / Arg) in GBS as it was raised in a previous control case study in India with GBS (Table 5). Here, we can see other associated diseases for ICAM-1 G241R polymorphisms studies. Differences can also be explained in part by statistical analysis methods (10,35).

**Table 5.** Associated diseases of 8 eligible studies for *ICAM-1* G241R polymorphisms analysis

| Diseases | Genotype | OR | p value | Population | Reference |
| --- | --- | --- | --- | --- | --- |
| Guillain-Barre Syndrome | 241Gly / Arg (GA) | 4.14 | <0.001 | Indian | (29) |
| Multiple Sclerosis | 241Gly / Arg (GA) | 0.64 | <0.200 | Polish Caucasian | (30) |
| Cancer | 241Gly / Arg (GA) | 2.03 | <0.010 | Asian-European-American | (31) |
| Cancer | 241Gly / Arg (GA) | 1.95 | <0.010 | European | (31) |
| Fuchs Uveitis | 241Gly / Arg (GA) | 3.3 | 0.012 | Italian | (32) |
| Schizophrenia | 241Gly / Arg (GA) | 1.14 | 0.771 | German Caucasian | (33) |
| Ischemic Stroke | 241Gly / Arg (GA) | 1.82 | 0.001 | ARIC Study (white) | (34) |
| Ischemic Stroke | 241Gly / Arg (GA) | 1.49 | 0.300 | ARIC Study (black) | (34) |
| Guillain-Barre Syndrome | 241Gly / Arg (GA) | 0.33 | 0.047 | Our study |  |

ARIC: Atherosclerosis Risk in Communities

The study number and sample size were limited, which may affect the reliability of the results. As well as, the differences may be explained by genetic diversities, different risk factors in life styles, and the exposure to different environmental factors. In conclusion, *ICAM-1* polymorphisms might be considered as potential genetic markers of GBS susceptibility after studies with larger sample size and further validation in ethnically different populations.

## ACKNOWLEDGEMENT

We are grateful to all the study participants. This work was supported financially by INBIOMEDIC, UTEC and Gen Lab of Peru.

